# Computational and experimental assessment of backbone templates for computational protein design

**DOI:** 10.1101/2021.06.23.449573

**Authors:** Frederikke Isa Marin, Kristoffer Enøe Johansson, Charlotte O’Shea, Kresten Lindorff-Larsen, Jakob Rahr Winther

**Author notes:** To whom correspondence may be addressed &. These authors contributed equally to this work (Frederikke Isa Marin, Kristoffer Enøe Johansson).

## Abstract

Computational protein design has taken big strides over the recent years, however, the tools available are still not at a state where a sequence can be designed to fold into a given protein structure at will and with high probability. We have here applied a recent release of Rosetta Design to redesign a set of structurally very similar proteins belonging to the Thioredoxin fold. We determined design success using a combination of a genetic screening tool to assay folding/stability in *E. coli* and selecting the best hits from this for further biochemical characterization. We have previously used this set of template proteins for redesign and found that success was highly dependent on template structure, a trait which was also found in this study. Nevertheless, state of the art design software is now able to predict the best template, most likely due to the introduction of the *cart_bonded* energy term. The template that led to the greatest fraction of successful designs was the same (a Thioredoxin from spinach) as that identified in our previous study. Our previously described redesign of Thioredoxin, which also used the spinach protein as template, however also performed well. In the present study, both these templates yielded proteins with compact folded structures, and enforces the conclusion that any design project must carefully consider different design templates. Fortunately, selecting designs using the *cart_bonded* energy term appears to correctly identify such templates.

## Introduction

In nature, proteins are specialized and highly efficient molecules and the functions of naturally occurring enzymes have already been recognized as useful in many products. Since natural protein sequences only explore a tiny fraction of the possible sequence space, it seems probable that proteins with improved properties or novel functions could be designed (1). Towards this goal, computational methods for full-sequence design have come a long way and it is today possible to design new sequences for proteins of known and novel folds (2–6).

Despite much progress and interest in the field, protein design is still hampered by low success rates, and medium- or high-throughput screening assays are increasingly used to handle the laborious task of testing many designs (5–8). Despite recent improvements in synthesis of DNA libraries (9,10), large scale studies of full sequence designs have mostly been limited to proteins <100 amino acids. In order to bridge the gap between the computational output and experimental screening, computational screens may be employed in addition to high-throughput screening technologies. However, there are few validated and generally applicable criteria by which to assess designs prior to experimental testing. Additionally, the designs will often fail to express, and the information gained from negative results is often ambiguous or uninstructive.

In this work, we report the test results of a complete protocol developed for full-sequence protein design. As a test case, we redesign, using recently developed methods, the Thioredoxin fold. This allows for a direct comparison with our previous work (11), primarily to address the issue of template selection in the design process. Thioredoxin (Trx) is a small protein (~110 residues), but still large enough to pose a significant challenge in design and larger than most *de novo* designs (4,6). Trx is globular with a compact fold and due to its evolutionary conservation throughout biology a wealth of homologs exists (12). Previous work showed that the success-rate of designs was highly dependent on subtle details of the template structure with one particular template outperforming the others (11). Here, we report improvements of the computational and experimental protocols and analyze the effect on the design workflow compared to our previous study. We found a substantial improvement in success rate by using a layered description in the computational protocol and by employing a folding sensor as a first experimental screen of the designs. *In vivo* folding sensors provide a powerful first screen for folded and stable designs and provide experimental information on successful and close-to-successful designs (5,13). Here, we employ such a sensor for which the sensitivity may easily be adjusted by changing the temperature (14). While we found that the template choice is still crucial for generating a reasonable success-rate, the software is now, interestingly, able to identify the best templates among the eight templates that we tested. We found that the template designability correlates with an energy term that reports on stress in covalent bond lengths and angles. The best template identified was the template (a Thioredoxin from spinach chloroplast, pdb 1fb0) that we previously identified, suggesting that there is something inherently designable about that structure. Interestingly, the runner-up in designability was the structure of a redesign of 1fb0 (pdb 5j7d), which is not more similar in structure to 1fb0 than the other targets.

## Experimental methods

### Computational design protocols

We tested two Rosetta design protocols based on the *FastDesign* procedure that iteratively optimizes sequence and structure, and the score function *REF15* (15) as implemented in *RosettaScripts* (16) GitHub SHA1 ce9cb339991a7e8ca1bc44efb2b2d8b0a3d557f8 of October 2018 (developer version). To evaluate the success rate as objectively as possible, both protocols are used automated and without subsequent ad hoc modifications of the protein sequences. Protocol 1 (P1) considers all amino acids, except Cys, at all positions whereas protocol 2 (P2) is restricted to a certain set of amino acids in regions of the structure with a similar degree of solvent exposure referred to as *layers*. Layered protocols are common and P2 is inspired by a previously published protocol (4). We used a three-layer protocol where only the amino acids VILMFYWGAP were allowed at core positions and DENQKRSTHPG at solvent exposed positions. The core was defined as having 5.2 or more side-chain neighbors and exposed positions with 1.9 or less neighbors, using the weighted neighbor count of the *LayerSelector* in *RosettaScripts*. In the boundary layer between core and exposed positions, we allowed all amino acids except MFHC. Both protocols had three additional restrictions; Conformational optimization of the backbone was restricted to a C_α_ RMSD of 1.5 Å from the template structures in order to retain the Thioredoxin fold, the number of unsatisfied hydrogen bonding donors and acceptors in the core was restricted to a maximum of 4, and the aromatic residues F, Y and H, were at all positions limited to a χ_2_ angle ranging from 70° to 110°, the range frequently observed in nature (4). All scripts, protocols and design output are available at https://github.com/KULL-Centre/papers/tree/master/2021/trx-redesign-marin-et-al.

Energy distribution are shown with kernel smoothing of bandwidth ~0.38 as determined by Scott’s rule (17). Sequences were aligned using ClustalW2 (18) and subsequently clustered with a neighbor joining algorithm with the same software. Resulting (phylogenetic) trees are visualized with Figtree (19). The branch lengths of the trees represent the distances between sequences.

### Experimental screening of designs in CPOP

Designed sequences for testing were custom synthesized and cloned into the CPOP folding sensor (14). Briefly, the CPOP system utilizes the enzyme orotate phosphoribosyl transferase (OPRTase), which is essential for growth of *E. coli* on medium lacking uracil. The sensor is based on a circularly permutated variant of OPRTase, with reduced stability. When designed sequences are inserted into this, misfolded designs will disrupt the structure of the enzyme and render the cells inviable on uracil-lacking medium whereas successful designs will enable growth. The severity of the folding defect can be gauged by assaying growth at different temperatures. Thus, CPOP provides a medium-throughput primary assay for the folding and stability of the designed proteins.

### Biophysical characterization

Designs that were able to complement growth at 37°C or higher were selected for further experimental characterization. For production of designs in *E. coli* outside the CPOP context, sequences were equipped with an N-terminal Met start codon, a C-terminal His_6_ tag, and expressed using a pJExpress441 expression vector (DNA2.0, Menlo Park, Calif., USA) in MC1061 cells. Designs showing significant amounts in the soluble fraction after sonication were further purified using Ni-NTA affinity chromatography. Designs with a solubility above 100 μM after purification were progressively tested for structural integrity as summarized in Table 2, initially using size exclusion chromatography (SEC) on a Superdex75 GL 10/300 analytical column. If the SEC showed a single peak of the expected retention time designs were further characterized by 1D proton NMR spectroscopy as described previously (14). Passing these tests, reversible two-state folding and stability was attempted using equilibrium unfolding with GuHCl with fluorescence excitation and emission at 280 and 360 nm, respectively (20).

## Results and discussion

### Computational design

The computational redesign of the Thioredoxin fold was based on eight backbone templates taken from the protein data bank (PDB). Seven of the templates were the same as in our previous study, however, the poorest performing template, 1dby, was replaced by the X-ray structure of a design from the same study, 5j7d (11). The latter was based on the best performing template, 1fb0, and could be considered a “second-generation” redesign of 1fb0. The templates were selected to have a gap-free alignment, minimal sequence identity (average 34%; Table S1) and maximal structural similarity (average C_α_ RMSD 1.2Å). We generated 120 computational designs for each backbone template and both protocols resulting in a total of 1920 sequences. The designs were denoted first by the protocol, then by PDB accession IDs of their template, and lastly by a rank according to their total Rosetta energy among the 120 designs (rank 1 having the lowest total energy). E.g., the best ranking design based on the 1fb0 template from P1 would be P1_1fb0_1.

As we have seen previously (11), the designed sequences are highly dependent on the choice of template (Figure 1 and Figure S1) even though these are very similar in terms of structure. The 120 designed sequences for a given template and protocol have an average of 39±3% and 40±3% pairwise identity for P1 and P2 respectively (Table S2). In comparison, designs, from the same template but different protocols, were only slightly more divergent with an average of 34±3% pairwise identity. On the other hand, designs are substantially more divergent when changing the template but not the protocol with an average of 26±3% and 28±3% (error estimates here and below correspond to standard deviation) pairwise identity between designs from different templates in P1 and P2 respectively (Table S2). Thus, we saw a larger effect on the sequence output from changing the template than from changing the protocol, reiterating the importance of template choice. Given different templates, the sequence diversity did not increase further by changing protocol also, since different templates and protocols resulted in average design identity of 24±3% (Table S2). In a phylogenetic analysis, the protocol effect is also strong enough to cluster sequences from the same protocol together in almost all cases (Figure S2), providing additional evidence for the template having a stronger effect than protocol on the sequence diversity.

**Figure 1.**
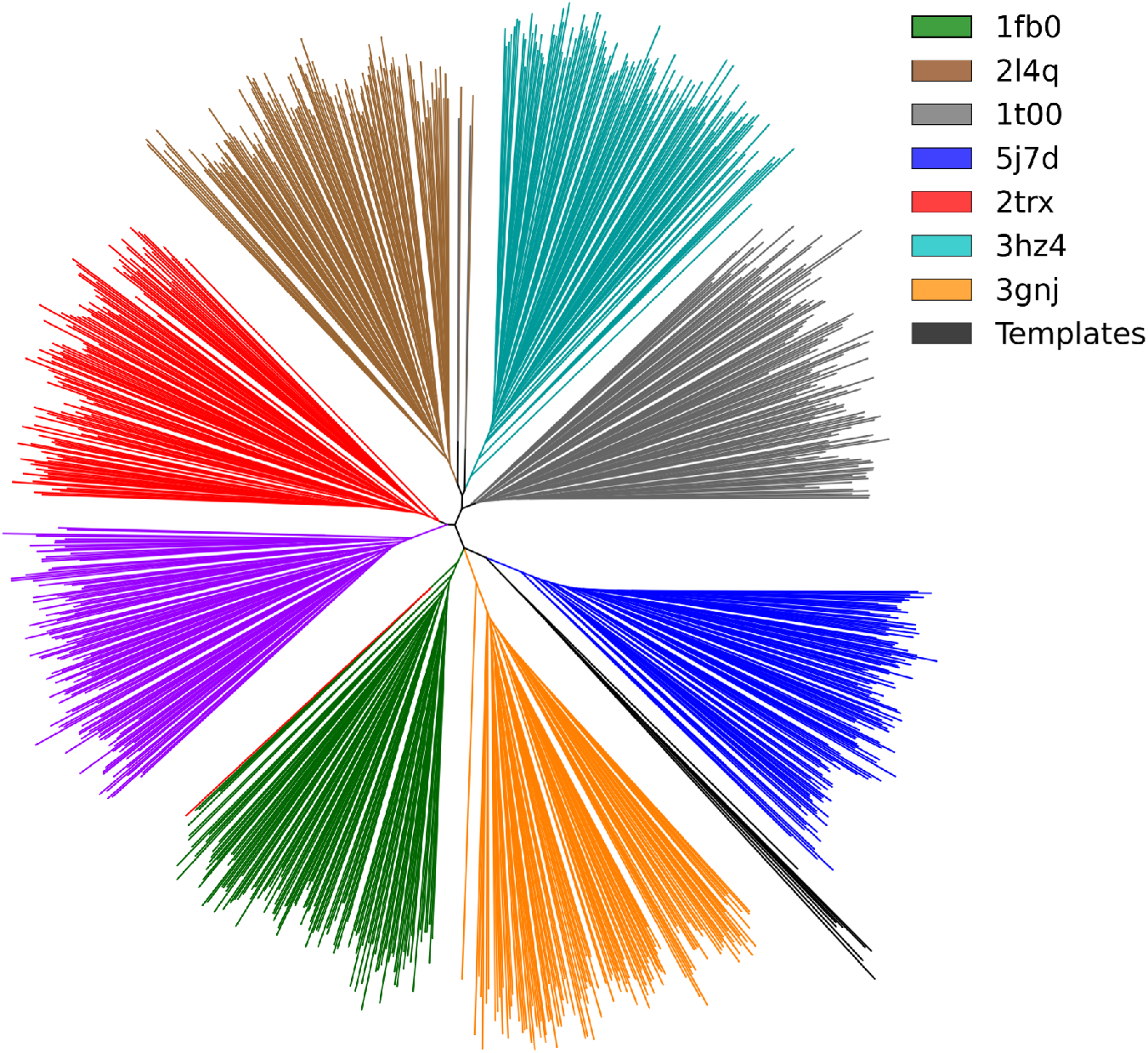
Phylogenetic tree of the 960 designed sequences from P2 together with the eight template (“wild-type”) sequences. Branches are colored according to the template used in the design, with template wild-type sequences in black. The lengths of the branches indicate the fraction of pairwise sequence identity. We highlight that the template sequences are closer to each other than to any of the designed sequences and that only very few of the sequences cluster with a different template.

While the template has a strong influence on the sequence diversity, this is not simply explained by the design recovering the template (wild-type) sequences. Thus, while the designed sequences resulting from a given template had an average pairwise identity of (P1) 39±3% and (P2) 40±3%, they only recaptured 25±4% and 22±4% of the wild-type sequences on average for P1 and P2, respectively (Table S3). This is also illustrated by the divergent branch of the wild-type sequences (Figure 1). The substantially lower recovery of the wild type sequence also indicates that the sequence bias that leads to clustering is not only towards the wild type but also towards something else that is dependent on the template structure and likely Rosetta protocol. We have previously seen that this bias is enhanced by the conformational relaxation and thus, that it becomes significant whether the iterative design protocol starts with optimization of conformation or sequence (11).

The sequence diversity was overall the same for the same template in the two protocols, but more divergent when compared to our previous study (~40% identity here versus 61% previously) (11). For different templates in the same protocol, the sequence diversity is similar to what we have seen previously (~27% identity here versus ~30% previously).

There are positions that are recaptured in a large fraction of the designs, most notably P61, G81 and G89 are recaptured in 80-100% of designs across all templates and protocols (Figure S3).

These positions are also conserved among native Thioredoxins which may indicate a role in structural stability (21), and for example in the spinach Trx structure (1fb0) glycine at positions 81 and 89 adopt positive ϕ backbone dihedral angles in line with the specific preferences of this amino acid (22). Our previous redesigns of the Thioredoxin fold, in general showed a higher degree reproduction of native residues, e.g. Val22 and Phe24 (11), which were only reproduced in ~50% of the design here.

### Experimental valuation of folding and stability of designs using the CPOP system

In our previous study (11), the frequency of designs that could be expressed in *E. coli* and purified with reasonable yield was 1-2 out of 48 (depending on experimental effort) and thus, we expected it to be necessary to screen at least this number of designs. In order to ease the experimental process, we employed the CPOP system that may both be used to screen and select for folding properties of designed proteins (14). CPOP is based on insertion of designed sequences into an intrinsically destabilized enzyme necessary of uracil biosynthesis in *E. coli*. Testing uracil requirement at different temperatures allows for estimation of folding competence of the inserted design. For CPOP testing, the four sequences with the best Rosetta energy scores were chosen from each template and from both protocols, resulting in 64 sequences in total. In terms of pairwise sequence identity, the selected sequences have a higher intra-template identity, but otherwise reflect the total set of designs well (Tables S4 and S5). On the other hand, this objective selection criteria enables us to compare the computational protocols and templates based on the screen. For each temperature, we categorize the level of complementation based on *E. coli* growth as either wild-type-like (green), significant but lower than wild-type (yellow) or insignificant (gray; Table 1 and Figure S4a).

**Table 1.**
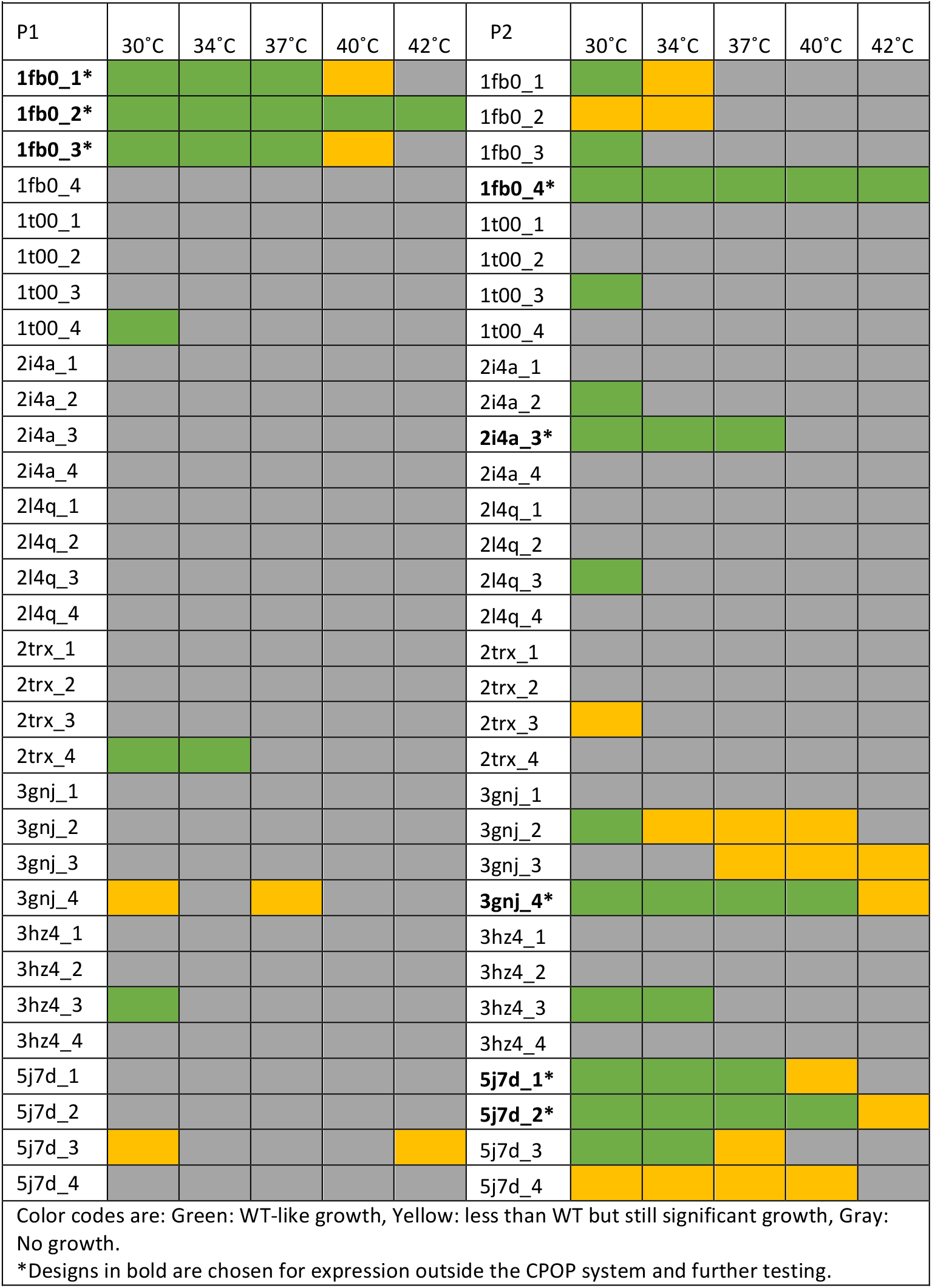
Growth of designs in the CPOP system at different temperatures.

**Table 2.**
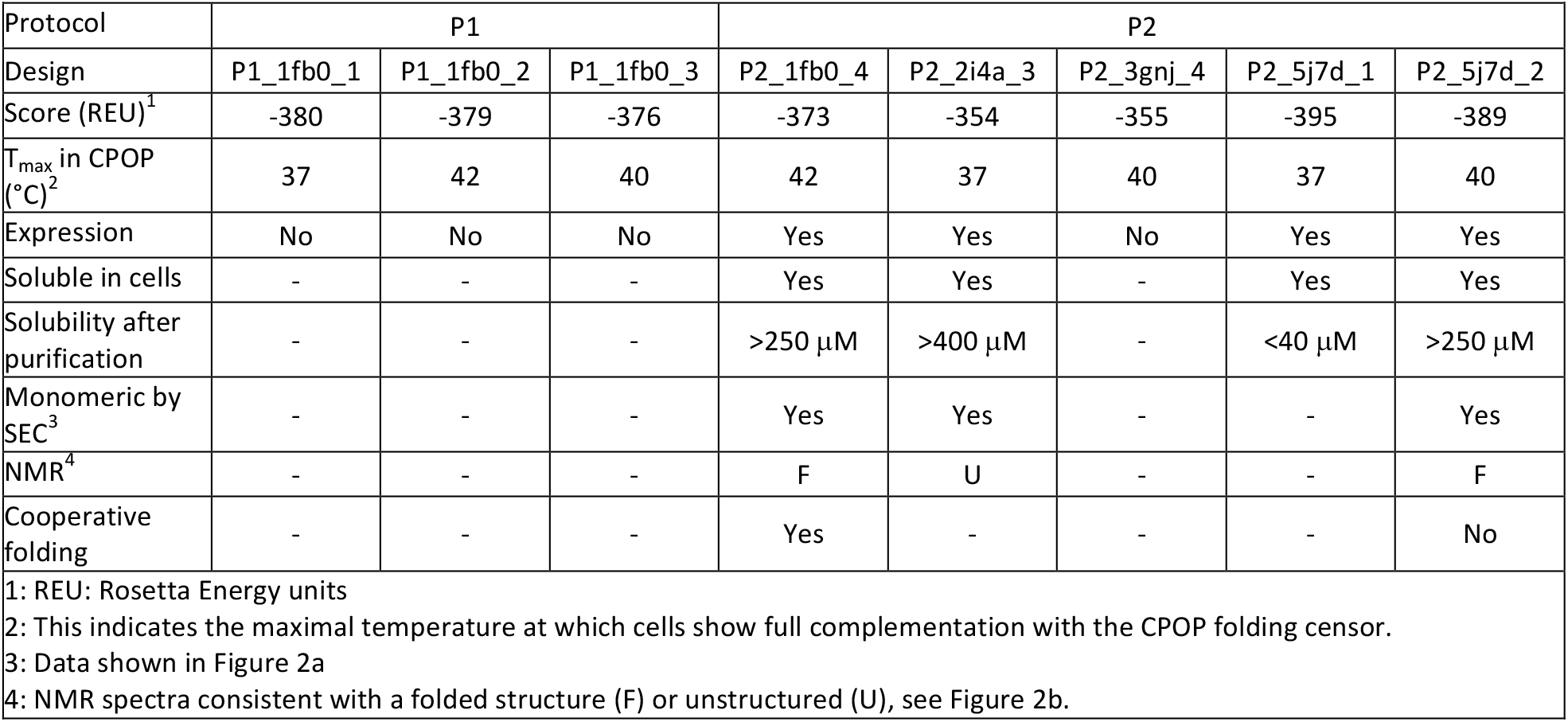
Summary of the characteristics of the eight designs scoring best in CPOP.

Despite small number of designs tested for each template the general trends of well-performing designs in P2 and certain templates are clear. What is not clear, for example, is if the decrease in complementation between P1_1fb0 and P2_1fb0 is an indication that 1fb0 produces better results in P1 than P2.

The layered protocol, P2, produced more designs with some level of complementation (17/32 or 53%) than did P1 (8/32 or 25%). The latter is comparable to our previous study where 26% (9/35) of designs showed any level of complementation (14). The major difference between the P1 and P2 protocols is that in P2 only certain types of amino acids are allowed in the core and another set on the surface. Thus, the set of all possible designed sequences from P2 are a subset of the sequences allowed in P1. Nevertheless, in the context of the Rosetta sequence optimization procedure, the diversity obtained in P2 is comparable to that in P1. Together with the results from the CPOP experiments this suggests that the restraints of P2 are highly relevant, since only a minor reduction of sequence diversity results in a substantial increase in CPOP performance.

A strong template effect was also observed in the CPOP screen with almost half of the complementing designs (12/25 or 48%) resulting from either 1fb0 or 5j7d (second-generation 1fb0 designs). Likewise, designs from these template structures generally complemented in CPOP to higher temperatures. Defining wild-type-like complementation at ≥37°C as “promising”, we found that 75% (6/8) of promising designs are from these two templates. The template 1fb0 was the most successful with 50% (4/8) of designs from both protocols screened to be promising. In comparison, in our previous study the designs based on 1fb0 that were tested in CPOP showed that only 7% (1/14) were promising with complementation ≥37°C (14).

Interestingly, the screen showed that the layered protocol, P2, was able to produce promising complementation ≥37°C with more templates than both P1 and our earlier protocol (11,14); in addition to 1fb0, P2 also generated promising designs based on 5j7d, 2i4a and 3gnj. All four tested designs from three templates (1t00, 2l4q and 3hz4), however, resulted in very poor complementation, which again highlights the strong template effect. In our previous work, the templates 2i4a and 3gnj resulted in the highest yields when expressed in *E. coli* but none of these designs could be purified (11) or complemented growth in the CPOP system (14). The observation that a few designs based on 2i4a and 3gnj here seemed closer to success in CPOP was thus consistent with the previous results, and suggests that the higher success-rate of the layered protocol is general and applies to all templates.

### Biophysical characterization

To test the performance of the designs independently of the CPOP system, we attempted to express and purify the eight promising designs that showed complementation in CPOP at ≥37°C (Table 2). Although three designs from P1 complemented to ≥37°C in the CPOP system, none of these could be expressed in *E. coli*. Expression success-rates may reflect stability but also other parameters that the computational design protocol does not consider. Our previous results show that complementation in the CPOP system correlates with yield in expression and solubility (14), however, the extent to which the circularly permutated OPRTase affects the folding, solubility and stability of the design has not been investigated in detail.

Three of the eight designs, P2_1fb0_4, P2_2i5a_3 and P2_5j7d_2, could be purified to a concentration required for further biophysical characterization. Although not a complete success, purified protein represents a minimum of design success from which further experimental data may be obtained in order to characterize or improve the design. Without purified protein, further experimental efforts to obtain a folded soluble protein are highly challenging, and the CPOP screening brings this success rate to 38% (3/8). Although we do not know how many of the 64 designs tested in CPOP could have been expressed and purified, this is comparable to some of the best reported design success rates for αβ-folds (6). Furthermore, even in the case that none designs could have been purified, the CPOP platform may also be used to select for rescue mutations (14).

Size exclusion chromatography (SEC) showed that all three designs that could be purified were monomeric with the expected elution volume (Figure 2A). As the proteins were all soluble to at least 250 μM, we recorded 1D NMR spectra to determine if they were consistent with folded structures. P2_1fb0_4 and P2_5j7d_2 showed a spectrum with upshifted methyl peaks and a good dispersion in the amide region of the spectrum consistent with a folded structure (Figure 2B) while P2_2i4a_3 appeared not to be well folded. We used chemical unfolding with guanidinium chloride to examine the stability of P2_1fb0_4 and P2_5j7d_2; however, while both appeared to fold reversibly, it was not possible to fit the data from P2_5j7d_2 to two-state folding and thus hampering determination of their thermodynamic stability. Interestingly, the template for P2_5j7d_2, itself a redesigned protein, also displayed non-two-state folding (11), a phenomenon not uncommon for designed proteins (3,23–25). P2_1fb0_4, on the other hand, could be fitted to two-state folding (20) with a ΔG_u_ of −20±3 kJ/mol (Figure 2C). We note that the fitted m-*value* (6.8±1.0 kJmol^-1^M^-1^) is about half the value expected from the size of Trx (29) and of that observed previously for *E. coli* Trx (30). This may be an indication of folding intermediates and together with a rather short base-line for the pretransition folded state, this gives some uncertainty to the stability determination.

**Figure 2.**
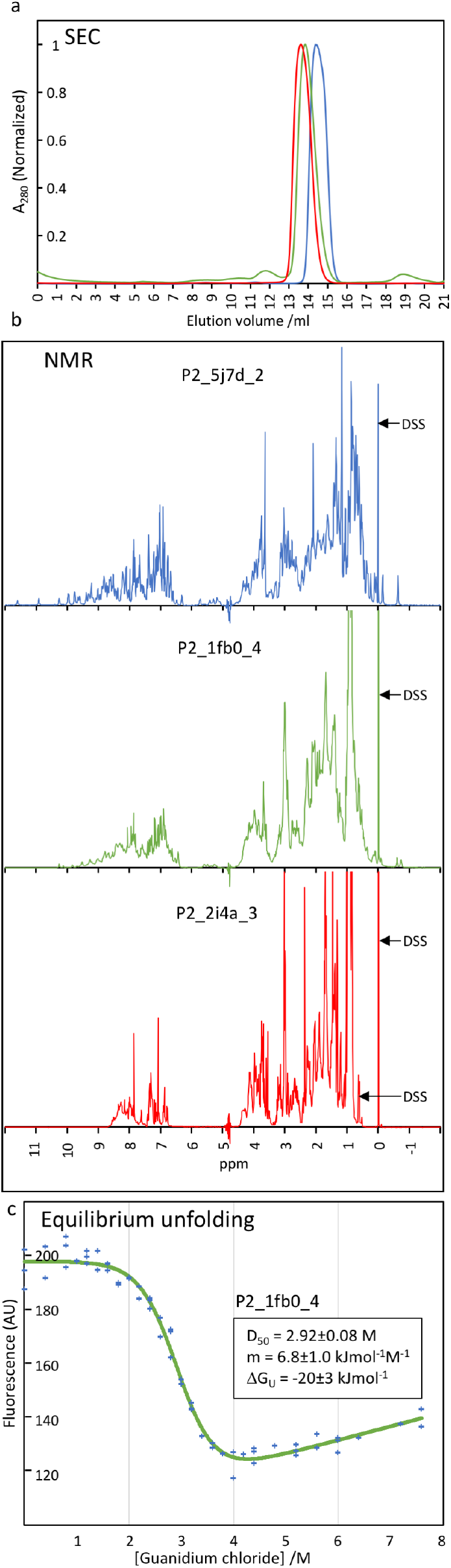
Characterization of successful protein designs. (a) Size exclusion chromatography of designs P2_5j7d_2 (blue), P2_1fb0_4 (green) and P2_2i4a_3 (grey) shows that all have an elution volume consistent with monomeric and compact structures. (b) 1H-NMR spectra of the designs from A indicated in the same color scheme as (a). Relative to P2_2i4a_3, the presence of up-shifted peaks in the methyl region (−1.0-1.0 ppm) as well as a dispersion in backbone amide region (6-11 ppm) in P2_5j7d_2 and P2_1fb0_4 are clear indications of well-defined folded structures. The sodium trimethylsilylpropanesulfonate (DSS) reference is indicated at 0 and 0.63 ppm. c) Chemical denaturation of P2_1fb0_4.

### Evaluation of design templates based on design energy

In our experimental analyses described above, we studied 64 sequences (32 per protocol)—evenly distributed over all templates—enabling us to assess possible differences in template designability. In an alignment of the eight sequences with lowest Rosetta energies (P2_5j7d_1-4 and P2_1fb0_1-4, combined) the level of conservation across designs is very low with only 7% completely conserved residues. Despite the improvements observed when using P2 instead of P1, our results reveal that the template still plays a substantial role in the success rate for the redesign task. In order to understand the effect of the backbone template, we examined the energy distributions of designs generated from each structural template (Figure 3 and Figure S5). In contrast to our previous work, Rosetta was here able to identify two successful designs, P2_5j7d_1 and P2_5j7d_2, with lowest energy among all designs from P2 independent of template. It is encouraging that simply selecting the two best ranking designs among all templates, would have resulted in a folded protein. Employing the CPOP and expressing the eight best designs from that, however, revealed the other most successful design, P2_1fb0_4, which would have been missed since design from 5j7d in general showed lower energies (Figure 3).

**Figure 3.**
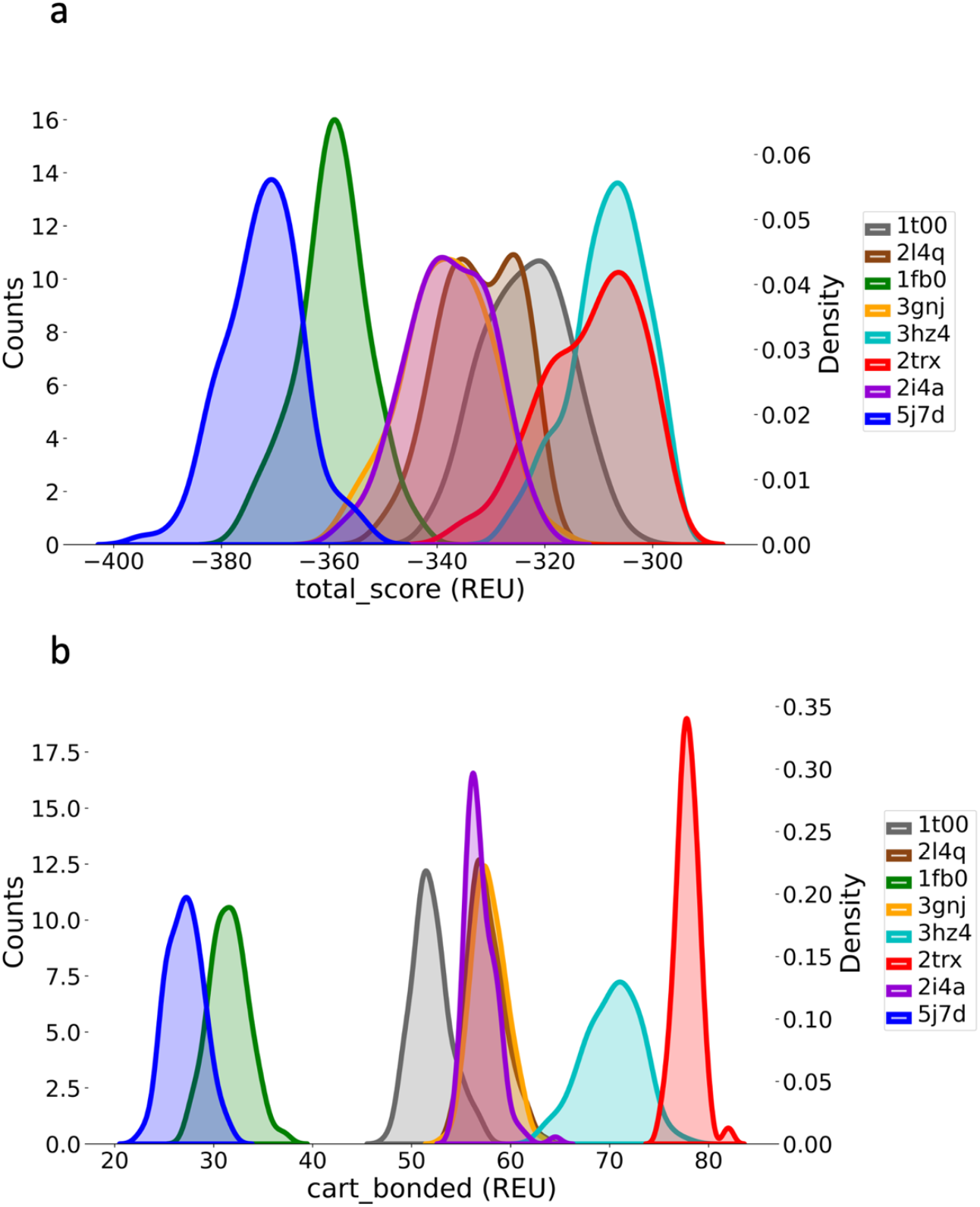
Rosetta energy distributions of all 120 designs per template for protocol P2. (a) Total Rosetta energy and (b) the cart_bonded energy term which is also part of the total energy summation. The cart_bonded term represents bond angles and lengths. All distributions have kernel smoothing for clarity.

Analysis of the individual components of the total energy revealed that the difference between design energy distributions mostly stems from the *cart_bonded* energy term (Figure 3b). This term was recently included in Rosetta with the *cartesian* protocol for conformational optimization that allows bond angles and lengths to deviate from their ideal values (26). The difference between the best, 5j7d, and worst templates, 3hz4 and 2trx, was, on average, 44 and 51 REU respectively. Exclusion of the *cart_bonded* energy term from the total energy reduced the separation of the total energy distributions for both protocols to an extent that it was difficult to discern any as being particularly favorable (Figure S6). This meant that *cart_bonded* energy term varied with template and therefore a property of the template structure. Further, it affects, or at least correlates, with the success of the resulting designs, and we did not find other correlating parameters (e.g. backbone angles) that distinguished the templates similarly to the *cart_bonded* term, nor which residues contributed to this. Conway et al. (26) found that conformational optimization in Cartesian space results in more accurate structure ranking and that the effect originates from allowing flexibility in the bond angles rather than bond lengths.

We also performed a conformational relaxation of the template structures with their wild-type sequences, which revealed the same pattern of the *cart_bonded* term (Figure S7), suggesting that the effect is a property of the template structure and sequence, and that it can be tested before designing new sequences. This might suggest that Rosetta underestimates the importance of bond angles in the sequence optimization and importantly, that this significantly affects the sequence outcome and design success even with conformational differences below 1 Å RMSD. Interestingly, a recent report of de novo design using extensive (human) sampling of backbone concluded that the cart_bonded energy term should be weighted four times higher (27). However, this knowledge seems not yet to have been propagated to newer versions of the Rosetta energy function.

In our previous study, we saw that very subtle changes in the design template could result in the reproduction of a buried Asp because the rigid side chain representation used in the design algorithm matched the steric environment of the template core very well. This was regardless of the fact that insertion of a buried Asp would come at an energy cost equivalent of shifting the *pKa* of the Asp to ≈7 (11). This highlights the great emphasis on steric core packing of the Rosetta energy function. In the work presented here, we have seen that a template with more relaxed bond lengths and angles results in a higher success rate and that this effect seemingly dominates over everything else tested here. This could indicate that the very precise core packing of Rosetta conserves bond stress present in the template. And since the precise packing match is only available for the stressed bond lengths and angles of the template, the most relaxed templates result in the highest success rate. What is surprising here, is that this seems to be the dominant effect on our success rate, though future prospective designs in other systems are needed to test this hypothesis.

### Evaluation of design protocols based on design energy

To evaluate the two design protocols used here, we looked at individual energy terms of Rosetta. In general, sampling constraints result in a higher energy because the algorithm is forced to avoid the low-energy regions that an un-constrained algorithm could sample. Thus, it is difficult to evaluate different protocols based on total energy whereas the individual energy terms that increase may indicate which energetic features are adjusted by the amino acid restrictions.

The score term *exposed hydrophobics* that quantifies the number and extent of exposure of hydrophobic side chains, is substantially higher in P1 than for P2 (Table S6 and S7 respectively). This term is not included in the total energy, but it reveals a tendency of unrestricted Rosetta to favor hydrophobic residues on the surface rather than penalizing it. This confirms the impact of avoiding hydrophobic residues at solvent exposed positions on design success rate. Solvent exposed hydrophobic residues may decrease solubility via unfavorable entropic water interactions, lead to unspecific interactions or even aggregation, or trigger the protein quality control system of the cell to degrade the protein. Rosetta is known to favor large side chains on the surface, driven by an inflated number of atomic interactions that are typically appear favorable to the energy function in the absence of explicit water (28).

## Conclusions

The present work assesses the state-of-the-art of protein re-design using eight backbone templates that are highly similar in terms of structure but not sequence. Using the CPOP *in vivo* folding sensor to screen 64 designs, we found that the strong template bias previously noted (11) still remains. While this initial screen clearly demonstrated the template preference, it also showed the strength of this biological assay as 3 of 8 best designs from this screen resulted in significant amounts of soluble protein, two of which resulted in folded protein as assessed by Far UV CD and NMR spectroscopy. Thus, we found that this screening approach was highly versatile and effective since it robustly identifies promising sequences.

While the overall design success rate is still dominated by the choice of backbone template, we also see significant improvements by using the layered P2 protocol that avoids hydrophobic residues at solvent exposed positions. Changing the template may increase the success rate from 25% to 75% for both protocols whereas changing the protocol improves from 25% to 53% based on CPOP complementation. We thus find that restricting the computational protocol in layers focuses the design on a more relevant subspace. As seen previously, designs of the different templates populate different regions of sequence space with significantly different success rates in both CPOP screening and biophysical characterization. A comparison of the structurally similar templates shows that Rosetta will, better than previously, reveal which templates are most designable. Our analysis suggests that the latter is related to the recent addition of the *cart_bonded* energy term and that this may be under-weighted in current versions of the Rosetta design energy function. The P2 protocol tested here raises the success rate of more templates to give useful results with the limited experimental effort of this study while only resulting in a minor tradeoff in output sequence diversity.

## Acknowledgements

This work was supported by Independent Research Fund Denmark to JRW, the Novo Nordisk Foundation Challenge programme PRISM centre (NNF18OC0033950; to KLL), and by cOpenNMR, Department of Biology, UCPH, an infrastructure granted from the Novo Nordisk Foundation (#NNF18OC0032996). The authors are grateful to Dr. Andreas Prestel for performing NMR spectroscopy.

## Data and code availability

All scripts, protocols and design output are available at https://github.com/KULL-Centre/papers/tree/master/2021/trx-redesign-marin-et-al.

