## Supplementary figures and tables for "Computational and experimental assessment of backbone templates for computational protein design"

(10 pages, 7 figures, 7 tables)

Supplementary figures:

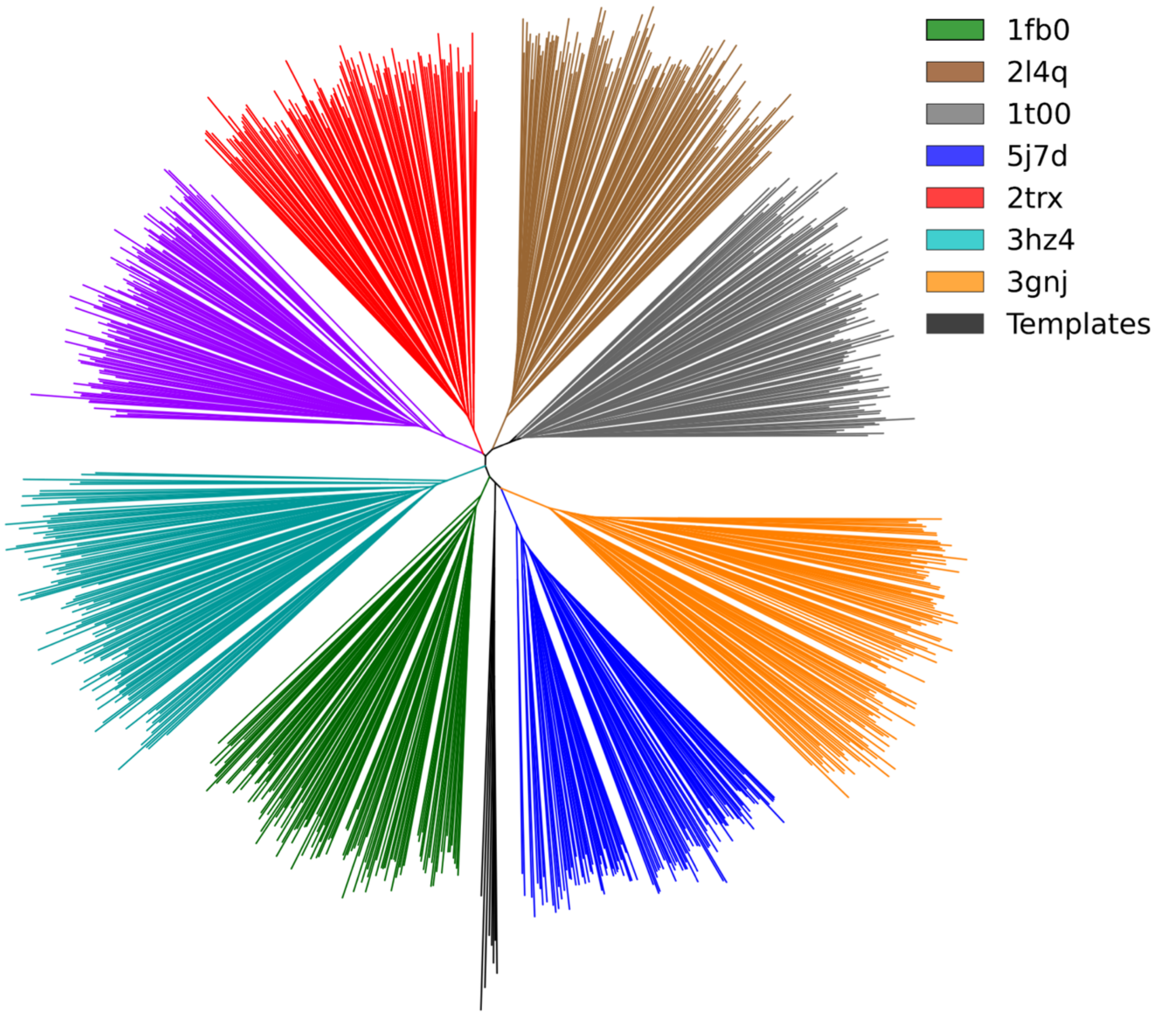


**Figure S1.** Phylogenetic tree of the 960 designed sequences from P1 and the 8 template sequences. Branches are colored according to their template, with wild-type template sequences in black. The lengths of the branches indicate the fraction of pairwise sequence identity i.e. the distance between the sequences. Sequences cluster according to their structural template but are divergent from the original template sequence (black).

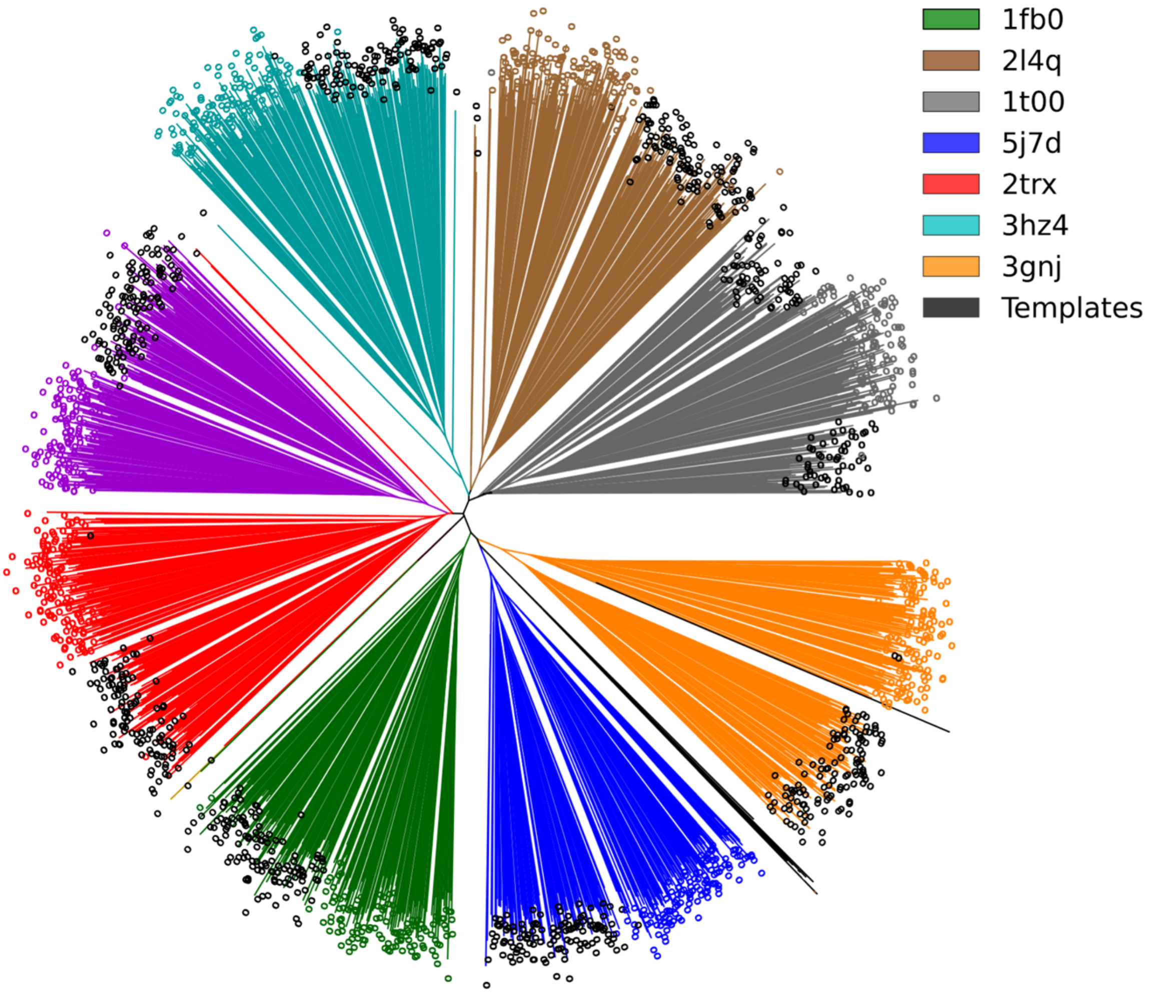


**Figure S2.** Phylogenetic tree of all 1920 designs from both protocols and the 8 wild-type template sequences. Branches are colored according to their template, with template sequences in black. Designs from P2 are marked with black circles and from P1 with colored circles.

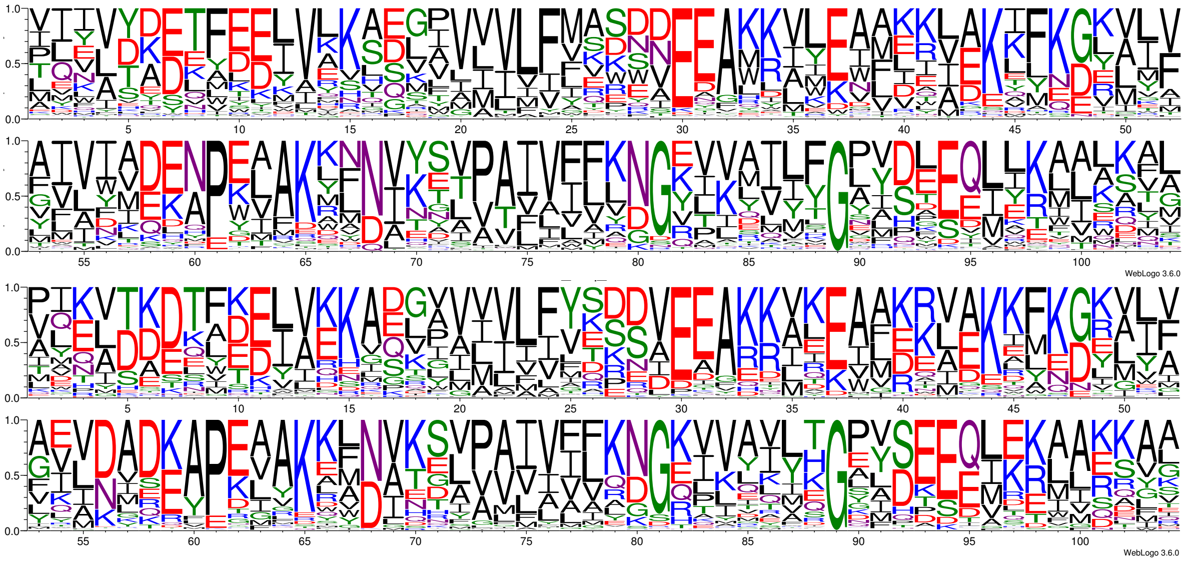


**Figure S3.** Sequence logo for all 960 designs generated by P1 (top) and P2 (bottom). Y-axis is frequency.

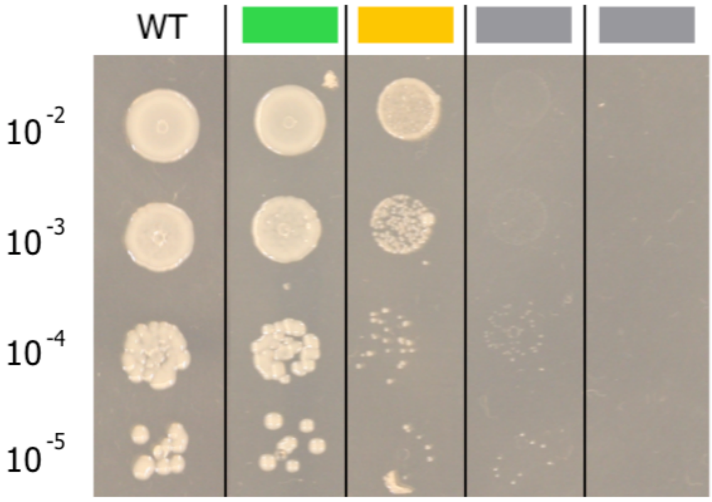


**Figure S4.** Experimental data. An example of a spot test. OD is normalized to 0.1. Colonies at the appropriate dilution (in this case 10-5) are compared to the WT. Complementation in the CPOP system (based on growth) is classified into three categories: Similar to WT (Green), less than WT but still significant growth (yellow), little or no growth (grey). The variants shown here for “similar to WT” and “less than WT” growth are P2_1fb0_4 and P2_1fb0_2 respectively. Both were grown at 30°C.

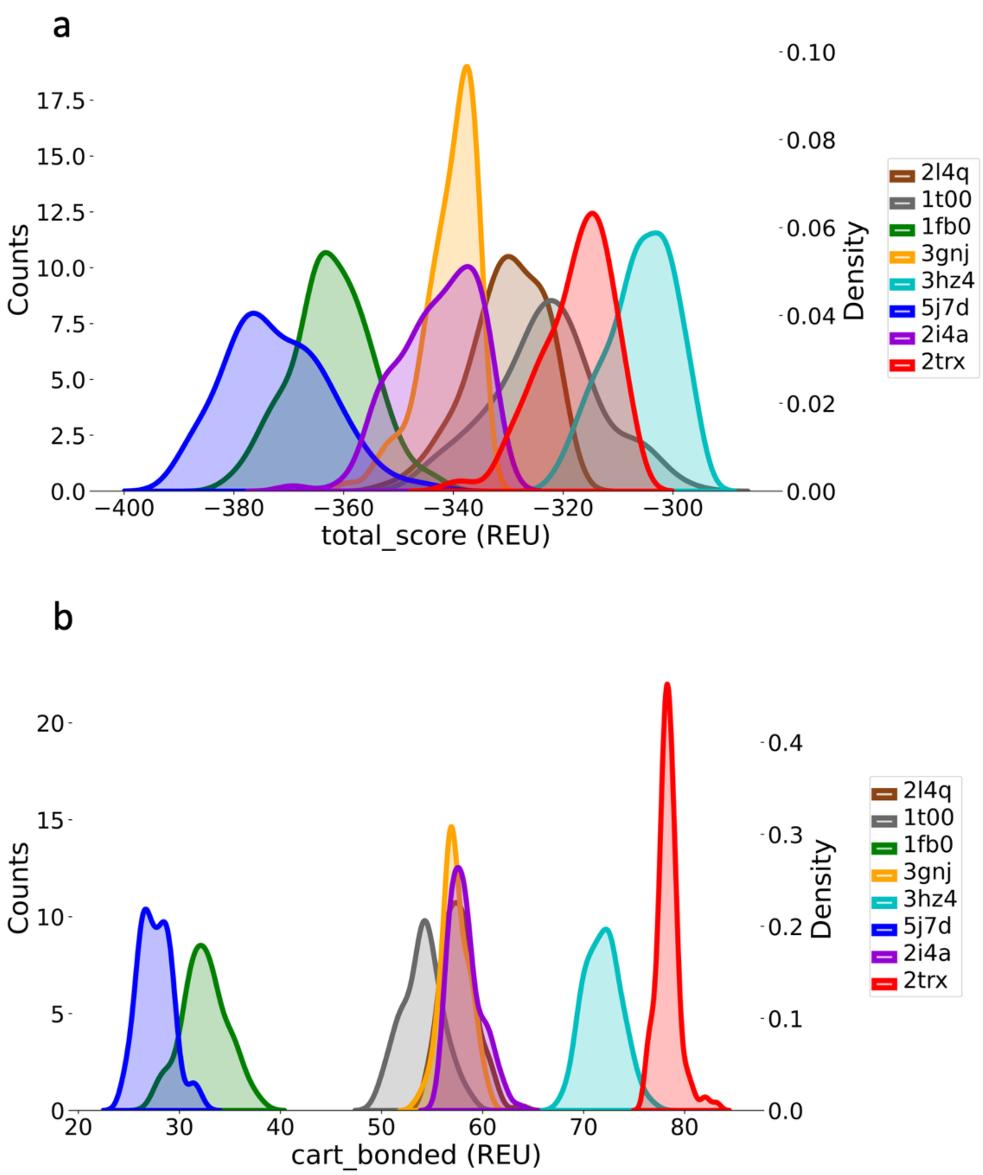


**Figure S5.** Rosetta energy distributions of all 120 designs per template for protocol P1. (a) Total energy and (b) the *cart_bonded* energy term which is also part of the summation in (a). The *cart_bonded* term represents bond angles and lengths. All distributions have kernel smoothing for clarity.

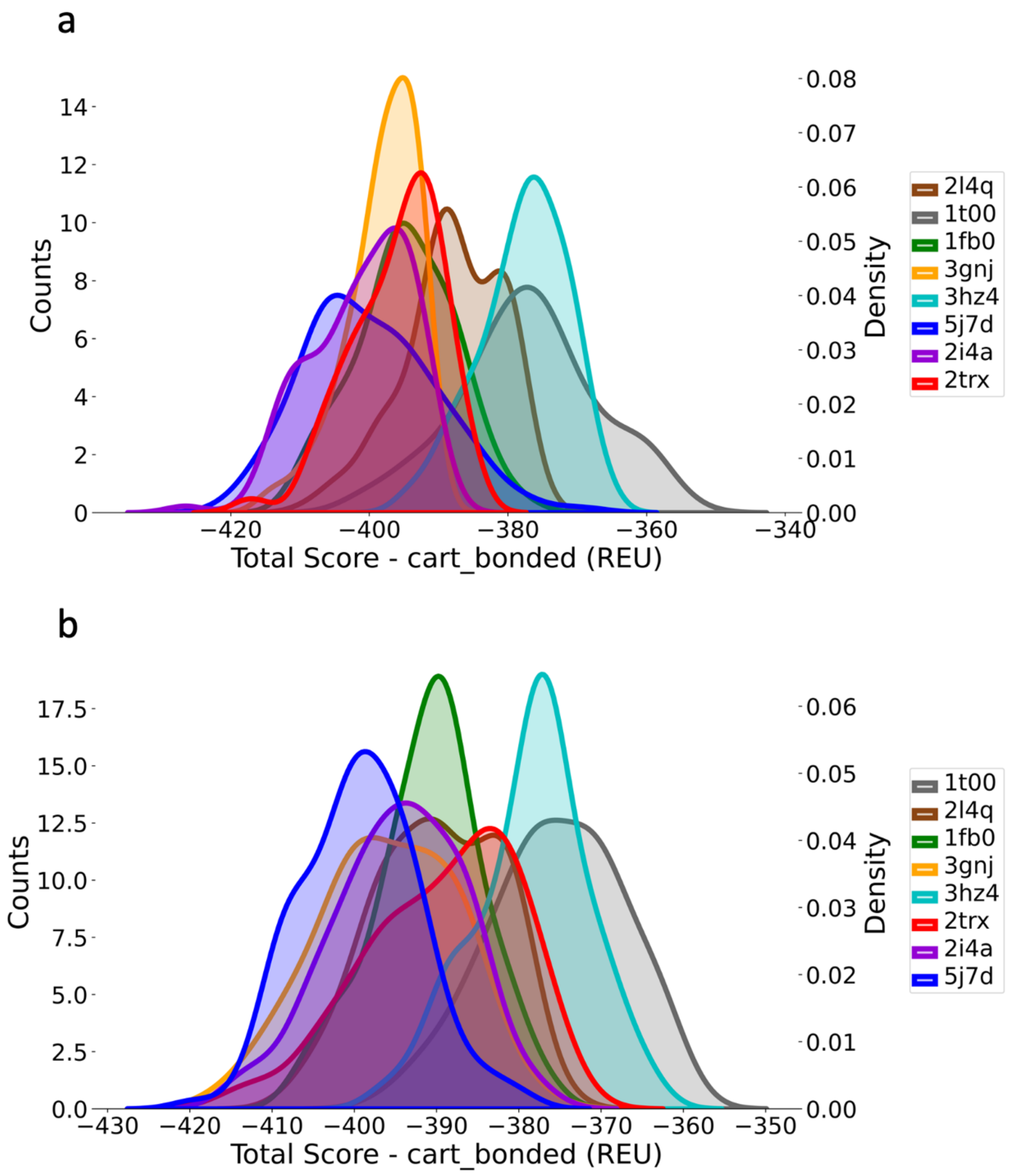


**Figure S6.** Total Rosetta energy of designs with the *cart_bonded* energy term subtracted (a) for designs from P1, (b) for designs from P2.

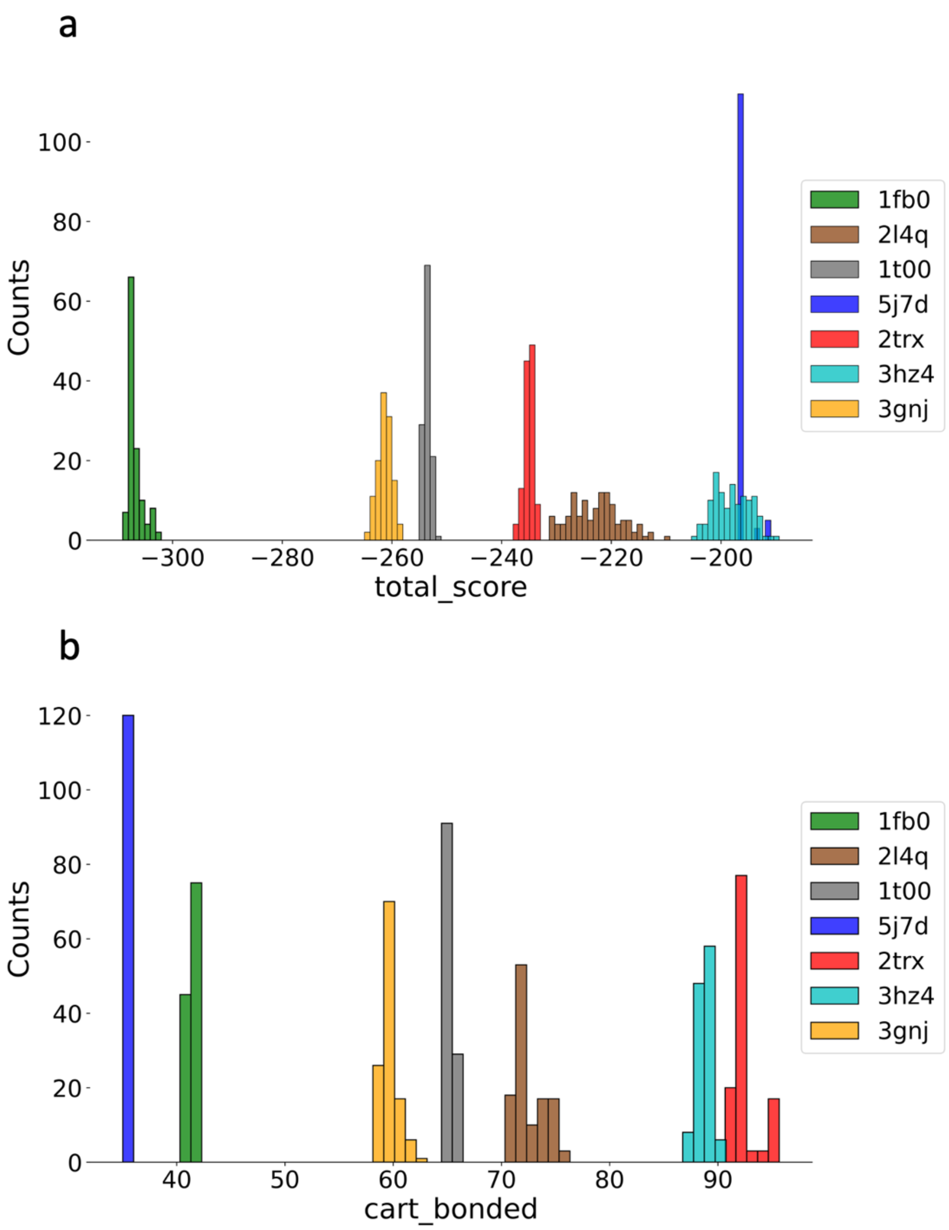


**Figure S7.** Rosetta energy of 120 parallel runs of *FastRelax* procedure for conformational optimization in Cartesian space for each of the structural templates used for design. (a) Total energy and (b) *cart_bonded* energy.

### Supplementary tables:

| | 1fb0 | 1t00 | 2i4a | 2l4q | 2trx | 3gnj | 3hz4 | 5j7d |
|---|---|---|---|---|---|---|---|---|
| 1fb0 | 100 | 44 | 42 | 41 | 48 | 21 | 30 | 47 |
| 1t00 | | 100 | 47 | 53 | 56 | 19 | 31 | 24 |
| 2i4a | | | 100 | 50 | 58 | 15 | 28 | 25 |
| 2l4q | | | | 100 | 52 | 24 | 32 | 28 |
| 2trx | | | | | 100 | 20 | 32 | 32 |
| 3gnj | | | | | | 100 | 21 | 17 |
| 3hz4 | | | | | | | 100 | 24 |
| 5j7d | | | | | | | | 100 |

**Table S1.** Pairwise sequence identity between WT template sequences in percent.

| | | P1 | | | | | | | | P2 | | | | | | | |
|---|---|---|---|---|---|---|---|---|---|---|---|---|---|---|---|---|---|
| | | 1fb0 | 1t00 | 2i4a | 2l4q | 2trx | 3gnj | 3hz4 | 5j7d | 1fb0 | 1t00 | 2i4a | 2l4q | 2trx | 3gnj | 3hz4 | 5j7d |
| P1 | 1fb0 | 39.9 | 27.7 | 28.6 | 25.9 | 28.0 | 24.8 | 25.2 | 30.4 | 35.2 | 26.9 | 27.7 | 25.0 | 26.6 | 24.3 | 24.9 | 28.3 |
| | 1t00 | | 37.6 | 29.1 | 28.3 | 29.5 | 22.0 | 24.1 | 25.6 | 26.7 | 31.9 | 26.7 | 26.8 | 26.4 | 21.9 | 22.8 | 23.9 |
| | 2i4a | | | 43.8 | 26.3 | 31.4 | 22.4 | 24.9 | 27.2 | 28.0 | 27.9 | 38.3 | 24.8 | 30.0 | 22.3 | 23.1 | 26.6 |
| | 2l4q | | | | 37.6 | 26.8 | 21.5 | 22.8 | 22.0 | 24.3 | 26.7 | 26.0 | 33.8 | 25.2 | 21.8 | 22.1 | 20.9 |
| | 2trx | | | | | 39.0 | 23.6 | 23.6 | 27.2 | 27.2 | 28.1 | 28.3 | 25.6 | 34.1 | 23.0 | 21.9 | 24.7 |
| | 3gnj | | | | | | 37.2 | 20.3 | 24.6 | 23.8 | 21.5 | 20.8 | 21.2 | 21.9 | 30.4 | 19.8 | 23.4 |
| | 3hz4 | | | | | | | 36.5 | 22.2 | 24.2 | 22.6 | 22.4 | 22.0 | 21.9 | 19.9 | 31.7 | 22.6 |
| | 5j7d | | | | | | | | 43.9 | 29.1 | 24.7 | 24.9 | 22.1 | 25.3 | 22.2 | 21.0 | 37.5 |
| P2 | 1fb0 | | | | | | | | | 41.0 | 30.8 | 32.1 | 27.5 | 30.1 | 27.4 | 27.5 | 32.1 |
| | 1t00 | | | | | | | | | | 37.9 | 31.9 | 30.7 | 31.1 | 24.6 | 26.2 | 27.0 |
| | 2i4a | | | | | | | | | | | 43.5 | 28.7 | 32.8 | 25.4 | 25.6 | 28.3 |
| | 2l4q | | | | | | | | | | | | 39.1 | 29.6 | 24.8 | 25.7 | 24.2 |
| | 2trx | | | | | | | | | | | | | 39.2 | 25.8 | 25.1 | 27.5 |
| | 3gnj | | | | | | | | | | | | | | 38.3 | 23.4 | 25.1 |
| | 3hz4 | | | | | | | | | | | | | | | 37.2 | 23.6 |
| | 5j7d | | | | | | | | | | | | | | | | 44.2 |

**Table S2.** Average pairwise sequence identity in percent between all 120 designs from each template and protocol.

| | | Wild type Sequence | | | | | | | |
|---|---|---|---|---|---|---|---|---|---|
| | | 1fb0 | 1t00 | 2i4a | 2l4q | 2trx | 3gnj | 3hz4 | 5j7d |
| P1 | 1fb0 | 29.1 | 19.1 | 21.7 | 20.5 | 21.7 | 17.9 | 19.1 | 31.2 |
| | 1t00 | 23.8 | 24.1 | 21.4 | 22.1 | 21.8 | 15.3 | 18.3 | 25.6 |
| | 2i4a | 19.3 | 19.6 | 26.0 | 20.7 | 23.6 | 13.9 | 16.1 | 23.1 |
| | 2l4q | 18.9 | 16.0 | 17.1 | 18.9 | 19.5 | 14.3 | 15.0 | 24.3 |
| | 2trx | 21.0 | 17.4 | 23.9 | 20.2 | 26.1 | 15.5 | 16.3 | 27.4 |
| | 3gnj | 18.0 | 13.4 | 15.7 | 14.9 | 17.0 | 22.0 | 13.7 | 25.0 |
| | 3hz4 | 16.9 | 19.1 | 16.9 | 17.3 | 18.8 | 13.2 | 19.5 | 20.7 |
| | 5j7d | 21.6 | 13.2 | 17.4 | 15.1 | 17.3 | 14.8 | 14.7 | 31.4 |
| P2 | 1fb0 | 24.8 | 17.6 | 21.0 | 17.8 | 20.5 | 16.3 | 18.1 | 30.6 |
| | 1t00 | 20.4 | 19.2 | 19.6 | 18.5 | 21.3 | 14.7 | 16.7 | 25.4 |
| | 2i4a | 20.0 | 18.6 | 23.5 | 19.0 | 23.1 | 13.9 | 16.3 | 23.0 |
| | 2l4q | 17.4 | 16.1 | 17.0 | 18.1 | 18.9 | 14.3 | 15.3 | 26.2 |
| | 2trx | 19.8 | 15.3 | 22.7 | 18.8 | 24.3 | 15.7 | 15.4 | 26.9 |
| | 3gnj | 17.5 | 12.3 | 16.3 | 14.6 | 18.0 | 18.9 | 15.4 | 21.6 |
| | 3hz4 | 17.5 | 17.9 | 16.3 | 16.7 | 17.9 | 13.9 | 17.7 | 23.3 |
| | 5j7d | 21.2 | 15.4 | 18.6 | 14.1 | 17.8 | 16.0 | 15.2 | 30.1 |

**Table S3.** Average pairwise sequence identity in percent between all 120 designs from one template and protocol and their corresponding WT template sequences.

| | | P1 | | | | | | | | P2 | | | | | | | |
|---|---|---|---|---|---|---|---|---|---|---|---|---|---|---|---|---|---|
| | | 1fb0 | 1t00 | 2i4a | 2l4q | 2trx | 3gnj | 3hz4 | 5j7d | 1fb0 | 1t00 | 2i4a | 2l4q | 2trx | 3gnj | 3hz4 | 5j7d |
| P1 | 1fb0 | 44.7 | 27.6 | 26.6 | 26.6 | 26.5 | 25.9 | 29.7 | 28.2 | 38.5 | 24.2 | 27.6 | 23.3 | 25.1 | 24.8 | 25.4 | 28.2 |
| | 1t00 | | 42.2 | 29.2 | 31.2 | 34.4 | 24.9 | 25.1 | 27.2 | 30.3 | 33.9 | 28.2 | 30.8 | 29.3 | 22.8 | 23.8 | 24.2 |
| | 2i4a | | | 52.5 | 26.3 | 32.0 | 22.9 | 28.4 | 28.1 | 26.9 | 26.9 | 41.4 | 25.1 | 30.2 | 24.9 | 24.2 | 28.8 |
| | 2l4q | | | | 44.7 | 28.8 | 23.3 | 26.4 | 21.4 | 24.4 | 27.2 | 25.1 | 37.2 | 27.8 | 22.8 | 22.3 | 18.9 |
| | 2trx | | | | | 45.3 | 23.9 | 26.7 | 28.2 | 29.9 | 30.8 | 26.6 | 26.4 | 37.6 | 23.8 | 20.9 | 24.1 |
| | 3gnj | | | | | | 38.8 | 24.7 | 24.9 | 24.6 | 18.6 | 20.0 | 20.9 | 22.1 | 28.1 | 22.4 | 23.1 |
| | 3hz4 | | | | | | | 39.8 | 24.9 | 26.4 | 23.5 | 24.9 | 23.9 | 24.4 | 20.6 | 29.4 | 25.4 |
| | 5j7d | | | | | | | | 56.3 | 27.9 | 23.9 | 25.1 | 21.1 | 27.8 | 21.2 | 21.4 | 37.1 |
| P2 | 1fb0 | | | | | | | | | 44.2 | 30.4 | 31.1 | 25.1 | 30.3 | 27.9 | 28.7 | 32.4 |
| | 1t00 | | | | | | | | | | 36.2 | 30.3 | 28.8 | 30.4 | 24.8 | 26.1 | 27.9 |
| | 2i4a | | | | | | | | | | | 47.7 | 29.4 | 31.5 | 25.9 | 24.6 | 28.4 |
| | 2l4q | | | | | | | | | | | | 39.0 | 31.9 | 22.8 | 25.8 | 21.3 |
| | 2trx | | | | | | | | | | | | | 40.5 | 26.6 | 24.6 | 25.0 |
| | 3gnj | | | | | | | | | | | | | | 44.7 | 28.4 | 25.4 |
| | 3hz4 | | | | | | | | | | | | | | | 38.3 | 23.9 |
| | 5j7d | | | | | | | | | | | | | | | | 51.0 |

**Table S4.** Average pairwise sequence identity in percent between 64 selected designs (the top four ranking for each template and each protocol).

| | | Wild type Sequence | | | | | | | |
|---|---|---|---|---|---|---|---|---|---|
| | | 1fb0 | 1t00 | 2i4a | 2l4q | 2trx | 3gnj | 3hz4 | 5j7d |
| P1 | 1fb0 | 30.5 | 20.8 | 22.8 | 21.2 | 21.8 | 18.2 | 21.5 | 32.0 |
| | 1t00 | 22.8 | 25.2 | 22.0 | 20.0 | 22.2 | 17.0 | 17.5 | 25.8 |
| | 2i4a | 18.5 | 20.5 | 25.2 | 21.2 | 24.0 | 13.2 | 16.8 | 22.2 |
| | 2l4q | 18.2 | 15.5 | 14.8 | 18.2 | 17.5 | 13.0 | 16.8 | 26.5 |
| | 2trx | 20.5 | 16.2 | 24.8 | 19.2 | 24.8 | 15.2 | 16.0 | 27.8 |
| | 3gnj | 19.0 | 12.5 | 15.8 | 14.2 | 15.8 | 24.0 | 14.5 | 24.0 |
| | 3hz4 | 17.0 | 18.5 | 19.0 | 18.2 | 18.5 | 15.0 | 22.0 | 21.8 |
| | 5j7d | 19.5 | 12.2 | 17.8 | 15.2 | 16.2 | 15.8 | 15.2 | 33.0 |
| P2 | 1fb0 | 29.2 | 19.5 | 22.0 | 20.0 | 23.2 | 16.8 | 18.0 | 34.5 |
| | 1t00 | 17.8 | 18.0 | 17.5 | 17.8 | 19.5 | 13.5 | 16.2 | 24.0 |
| | 2i4a | 20.0 | 20.0 | 22.5 | 18.2 | 25.8 | 12.5 | 15.2 | 22.0 |
| | 2l4q | 16.2 | 15.0 | 16.2 | 17.5 | 19.2 | 14.8 | 17.2 | 24.8 |
| | 2trx | 20.5 | 16.5 | 24.5 | 19.5 | 27.2 | 15.8 | 14.8 | 28.0 |
| | 3gnj | 15.0 | 11.5 | 16.5 | 13.8 | 18.0 | 17.5 | 15.8 | 20.5 |
| | 3hz4 | 17.5 | 14.5 | 17.0 | 13.8 | 18.0 | 16.2 | 15.8 | 25.0 |
| | 5j7d | 18.0 | 13.8 | 16.2 | 12.2 | 15.2 | 12.8 | 15.2 | 26.2 |

**Table S5.** Average pairwise sequence identity in percent between top 4 designs from each template and protocol and their corresponding wild-type template sequence.

| Score term | 1fb0_P1 | 1t00_P1 | 2i4a_P1 | 2l4q_P1 | 2trx_P1 | 3gnj_P1 | 3hz4_P1 | 5j7d_P1 |
|---|---|---|---|---|---|---|---|---|
| **Total score (REU)** | -362.3 | -323.03 | -342.82 | -330.3 | -317.65 | -340.75 | -305.84 | -372.23 |
| **cart_bonded** | 32.5 | 54.03 | 58.33 | 57.65 | 78.37 | 57.29 | 71.89 | 27.62 |
| **fa_atr** | -610.03 | -613.0 | -612.4 | -603.14 | -619.04 | -634.55 | -613.79 | -625.88 |
| **fa_rep** | 77.48 | 75.66 | 76.08 | 67.84 | 76.27 | 74.27 | 72.63 | 72.12 |
| **fa_intra_rep** | 1.3 | 1.32 | 1.24 | 1.32 | 1.33 | 1.41 | 1.28 | 1.32 |
| **fa_sol** | 281.18 | 292.18 | 305.0 | 272.01 | 282.53 | 303.27 | 302.04 | 297.66 |
| **lk_ball_wtd** | -11.13 | -12.0 | -11.48 | -11.31 | -12.4 | -11.22 | -11.59 | -11.78 |
| **fa_intra_sol_xover4** | 17.89 | 17.96 | 16.99 | 17.49 | 18.44 | 18.87 | 18.91 | 18.08 |
| **fa_elec** | -181.13 | -179.44 | -198.91 | -180.92 | -176.51 | -190.11 | -188.51 | -182.52 |
| **hbond_lr_bb** | -26.85 | -27.96 | -29.8 | -28.01 | -28.53 | -30.06 | -27.09 | -23.93 |
| **hbond _sr_bb** | -37.49 | -34.99 | -35.81 | -37.7 | -34.38 | -41.26 | -36.27 | -44.01 |
| **hbond_bb_sc** | -12.43 | -14.09 | -20.46 | -11.99 | -14.69 | -12.1 | -14.4 | -15.72 |
| **hbond_sc** | -14.67 | -13.38 | -11.97 | -12.42 | -13.4 | -12.75 | -14.75 | -13.6 |
| **rama_prepro** | -15.68 | -14.88 | -20.87 | -16.15 | -18.56 | -11.46 | -12.92 | -18.2 |
| **p_aa_pp** | -27.82 | -26.0 | -30.12 | -27.1 | -29.77 | -28.09 | -24.05 | -28.35 |
| **fa_dun** | 109.54 | 117.08 | 111.27 | 113.53 | 119.41 | 117.32 | 117.81 | 114.57 |
| **omega** | 2.29 | 5.75 | 4.55 | 12.21 | 2.83 | 8.32 | 5.66 | 5.56 |
| **yhh_planarity** | 0.06 | 0.09 | 0.11 | 0.05 | 0.05 | 0.04 | 0.03 | 0.03 |
| **ref** | 52.68 | 48.65 | 55.42 | 56.33 | 50.38 | 50.05 | 47.27 | 54.81 |
| **Additional score terms not included in the REU** | | | | | | | | |
| **Unsatisfied hbonds core** | 2.84 | 2.82 | 3.03 | 3.63 | 2.77 | 2.1 | 2.62 | 2.41 |
| **Packing holes** | -1.36 | -0.97 | -1.72 | -1.31 | -1.37 | -1.71 | -1.63 | -0.97 |
| **RMSD to WT** | 0.87 | 0.88 | 0.98 | 0.89 | 0.87 | 0.87 | 0.89 | 0.95 |
| **Recapture of WT seq** | 0.29 | 0.24 | 0.26 | 0.19 | 0.26 | 0.22 | 0.22 | 0.14 |
| **Exposed hydrophobics** | 1064.14 | 1148.12 | 1026.12 | 1170.48 | 1215.57 | 1182.97 | 1055.95 | 1204.37 |
| **Packing statistics** | 0.67 | 0.66 | 0.69 | 0.66 | 0.7 | 0.71 | 0.69 | 0.68 |

**Table S6.** Average values of energy term per template for the P1 protocol.

| Score term | 1fb0_P2 | 1t00_P2 | 2i4a_P2 | 2l4q_P2 | 2trx_P2 | 3gnj_P2 | 3hz4_P2 | 5j7d_P2 |
|---|---|---|---|---|---|---|---|---|
| **Total score (REU)** | -359.41 | -323.39 | -337.64 | -332.17 | -311.4 | -338.09 | -308.21 | -372.2 |
| **cart_bonded** | 31.56 | 51.98 | 56.98 | 57.59 | 77.88 | 57.86 | 70.27 | 27.11 |
| **fa_atr** | -590.29 | -591.05 | -581.42 | -582.4 | -597.04 | -603.02 | -593.81 | -598.61 |
| **fa_rep** | 73.82 | 72.47 | 70.4 | 64.62 | 72.97 | 69.57 | 71.34 | 65.67 |
| **fa_intra_rep** | 1.17 | 1.14 | 1.09 | 1.17 | 1.23 | 1.18 | 1.14 | 1.17 |
| **fa_sol** | 313.34 | 324.88 | 327.42 | 309.15 | 315.06 | 331.46 | 330.39 | 335.89 |
| **lk_ball_wtd** | -9.7 | -9.28 | -10.81 | -9.89 | -10.61 | -8.51 | -9.2 | -10.24 |
| **fa_intra_sol_xover4** | 18.97 | 18.52 | 17.51 | 18.49 | 19.53 | 19.61 | 19.49 | 20.52 |
| **fa_elec** | -206.96 | -210.05 | -218.82 | -211.23 | -205.63 | -214.99 | -216.41 | -216.28 |
| **hbond_lr_bb** | -26.73 | -28.52 | -30.25 | -28.3 | -29.21 | -29.57 | -27.79 | -23.72 |
| **hbond_sr_bb** | -38.36 | -35.92 | -35.74 | -38.55 | -34.5 | -41.87 | -36.27 | -43.77 |
| **hbond_bb_sc** | -11.59 | -14.26 | -21.09 | -13.09 | -17.27 | -14.19 | -15.67 | -14.51 |
| **hbond_sc** | -18.96 | -19.93 | -18.07 | -18.24 | -17.85 | -17.63 | -20.18 | -22.04 |
| **rama_prepro** | -16.1 | -13.34 | -19.13 | -15.89 | -17.73 | -11.0 | -11.68 | -17.7 |
| **p_aa_pp** | -25.71 | -22.39 | -27.31 | -25.37 | -27.13 | -26.27 | -22.58 | -26.35 |
| **fa_dun** | 117.0 | 117.31 | 114.03 | 117.54 | 125.09 | 121.78 | 122.82 | 122.82 |
| **omega** | 2.11 | 5.85 | 3.95 | 11.94 | 3.12 | 7.37 | 6.04 | 4.48 |
| **yhh_planarity** | 0.05 | 0.09 | 0.11 | 0.04 | 0.07 | 0.04 | 0.04 | 0.03 |
| **ref** | 26.98 | 29.09 | 33.51 | 30.25 | 30.64 | 20.12 | 23.86 | 23.33 |
| **Additional score terms not included in the REU** | | | | | | | | |
| **Unsatisfied hbonds core** | 2.77 | 3.22 | 2.73 | 3.52 | 3.1 | 2.54 | 3.0 | 2.59 |
| **Packing holes** | -1.44 | -1.39 | -1.72 | -1.65 | -1.4 | -1.81 | -1.75 | -1.25 |
| **RMSD to WT** | 1.05 | 1.15 | 1.15 | 1.09 | 1.11 | 1.09 | 1.19 | 1.14 |
| **Recapture of WT seq** | 0.25 | 0.19 | 0.24 | 0.18 | 0.24 | 0.19 | 0.19 | 0.16 |
| **Exposed hydrophobics** | 485.05 | 543.44 | 470.3 | 598.75 | 644.28 | 500.49 | 459.82 | 508.57 |
| **Packing statistics** | 0.64 | 0.63 | 0.64 | 0.63 | 0.66 | 0.68 | 0.66 | 0.64 |

**Table S7.** Average values of energy term per template for the P2 protocol.
